# High-resolution mapping of the MCT uncovers late blastomere commitment and DREAM/MuvB-mediated restriction of blastomere plasticity

**DOI:** 10.64898/2026.11.13.766562

**Authors:** Julien Lambert, Sophie Jarriault

## Abstract

Blastomeres are thought to lose multipotency rather sharply during early embryonic development in *C. elegans*, but the precise timing and regulation of this transition remain incompletely characterised. Here, we present a refined cell-fate challenge assay enabling high-resolution mapping of the multipotency-to-commitment transition (MCT). By combining precise heat-shock timing, 2-cell embryo synchronisation and single nuclei quantification, our method can detect subtle differences in blastomere plasticity across genotypes and developmental stages. Using this approach, we find that the MCT is more gradual than previously appreciated as blastomeres remain competent to adopt the body-wall muscle fate until at least the late 16E stage. We further show that the DREAM/MuvB complex core component LIN-54 restricts blastomere multipotency as does the H3K9 methyltransferase MET-2. Together, our work provides a sensitive single-cell assay for assessing developmental plasticity, reveals an extended timeline for the MCT and suggests that LIN-54/DREAM and MET-2 cooperate to promote blastomere commitment.

## Results

### Refined high-resolution cell-fate challenge assay

To test factors, such as *lin-15A* and its genetic interactors (Becker et al. 2026), that may impact blastomere plasticity or multipotency, we developed a method that measures Multipotency-to-Commitment Transition (MCT) timing with high temporal precision and sensitivity. This method relies on ubiquitous, heat-shock–induced expression of the muscle-determining factor HLH-1/MyoD in strains expressing nuclear NLS::GFP tagged under the control of a body-wall muscle (BWM)–specific promoter (Stringham et al. 1992) In such strains, heat-shock (hs) until the 8E stage, previously defined as the MCT, leads to most cells to adopt a muscle fate (Fukushige and Krause 2005; Spickard et al. 2018; Rothman and Jarriault 2019). The cells that differentiate into the BWM identity—whether naturally or through transdetermination— display a strong nuclear GFP signal. Given the robustness of this signal, we hypothesised that it would be possible to quantify with precision the number of cells differentiating into BWM, similar to the approach used by (Gilleard and McGhee 2001).

Timed hs are key to determine when is the latest developmental stage where over-expressed *hlh-1* leads to transdeterminations. To heat-shock embryos at defined developmental stages, we tested several synchronisation methods and found that the most precise approach was to dissect gravid mothers in M9 using sterile needles and a mouth pipette for embryo selection (Fig. 1B). We specifically collected 2-cell stage embryos (40–75 min post-fertilisation). To obtain enough embryos per experiment (10–20 embryos), several gravid mothers were dissected simultaneously. Selected embryos were transferred to individual wells of multi-well microscopy slides (mw-slides) in 8 µL of M9 (Fig. 1B). After loading, slides were sealed with melted VALAP (see Materials and Methods) and immersed in a 20 °C water bath until the desired hs time (200, 250, 280, 330, or 370 min post-fertilisation in this study).

**Figure 1.**
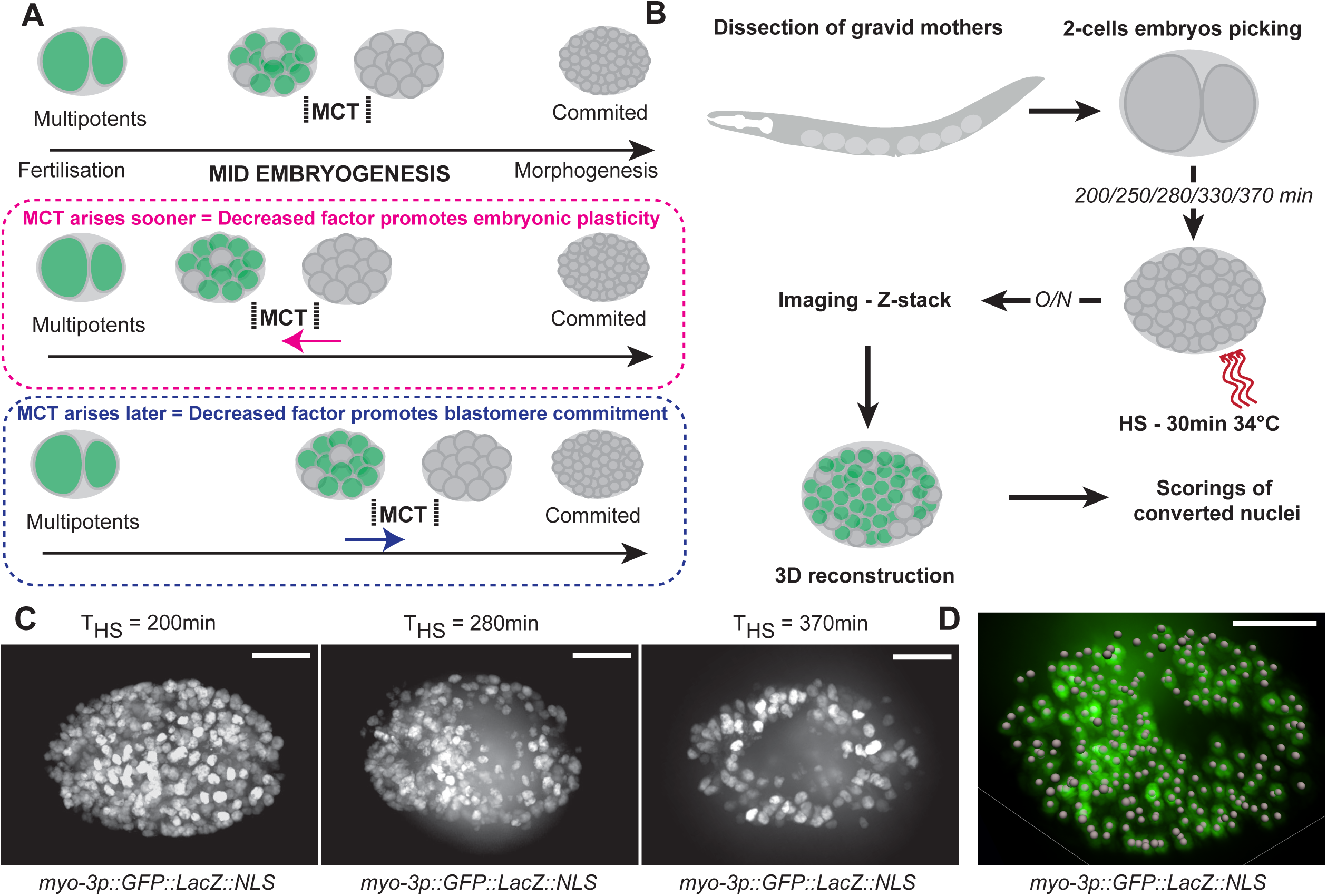
Experimental set-up of the high-sensitivity cell-fate challenge assay in *C. elegans* embryos. **A.** Top panel : Ubiquitous overexpression of cell fate determinants has been used to determine the timing of the MCT in *C. elegans*. For instance, exposure of blastomeres to the HLH-1/MYOD muscle determinant prior to the MCT leads to many of them transdetermining and their descendants adopting a muscle fate, while no or little transdetermination are observed with exposure after the MCT (green, stages at which blastomeres exhibit multipotency). Middle panel : MCT occurs sooner - Expected impact on the timing of the MCT upon the loss of a factor that would promote blastomere plasticity/multipotency ; Lower panel : MCT occurs later - Expected impact upon loss of a factor that would promote blastomere commitment/differentiation. **B.** Experimental set-up for the cell fate challenge assays. 2-cell embryos (45-70min post-fertilisation, or p-f) are picked from dissected gravid mothers, placed on multi-welled slides until the desired timing, heat-shocked, and imaged after over/night incubation. 3D-reconstruction allows to quantify both the total number of nuclei and the number of nuclei that are converted by the cell-fate challenge. **C.** Representative images (Max. projections) of embryos carrying *hsp-16.41p::hlh-1 cDNA* and *myo-3p::GFP::LacZ::NLS* transgenes, heat-shocked at 200 / 280 / 370min p-f, after overnight incubation. Left panel, at 200min, almost all blastomeres can be converted to BWM, while this response is lower at 280min (middle panel), and almost null at 370min (only the natural muscle-lineage being positive – right panel). Scalebars = 10 µm. **D.** Representative image of a 3D-rendering from an embryo carrying *hsp-16.41p::hlh-1 cDNA* and *myo-3p::GFP::LacZ::NLS* transgenes, heat-shocked at 280min (late 8E stage). The grey spheres indicate all the nuclei recognised and scored after segmentation using Imaris. Scalebar = 10 µm

At the desired time, mw-slides were immersed in a 34 °C water bath for 30 minutes to induce expression of HLH-1/MyoD under the control of the *hsp-16.41p* promoter (Fukushige and Krause 2005) (Fig. 1B). After heat shock, slides were returned to a 20 °C water bath for overnight incubation, allowing blastomeres—successfully or unsuccessfully challenged—to differentiate into their terminal fates. The following day, embryos were imaged and analysed using full-embryo 3D reconstruction (Fig. 1B,C,D). Importantly, under these conditions embryos, even those fully converted, remained alive, and did not hatch after overnight incubation.

This experimental procedure and its sensitivity were validated by the different levels of response observed when embryos were heat-shocked at distinct timings (Fig. 1C). For instance, embryos heat-shocked before the MCT, at 200 min post-fertilisation (4E stage), showed complete sensitivity to HLH-1 exposure, with almost all blastomeres differentiating into BWM *myo-3p::GFP*–positive cells. In contrast, response was much lower for 370 min (late 16E stage) hs, while embryos heat-shocked at 280 min (late 8E stage) displayed an intermediate phenotype.

### Timing corrections for strains with different embryonic developmental speeds

In *C. elegans*, numerous mutants exhibit reduced embryonic developmental speed. A well-known example is the mitochondrial *clk-1* mutants (Wong et al. 1995; Vajo et al. 1999). However, this effect is not limited to metabolic mutants. Notably, losses of LIN-54 or LIN-15B was previously shown to slow embryonic development (Costello and Petrella 2019). Therefore, to hs genotypes with different developmental speeds at equivalent developmental times, we applied timing corrections.

To do so, a reference strain that expresses a nuclear GFP under the control of the *end-1p* promoter in the E lineage (*teIs46* integrated array) was used (Fig. 2). In this strain, we measured the duration between fertilisation and the transition from the 4E to the 8E stage. In parallel (using different embryos to avoid photo-toxicity), we measured the duration between fertilisation and the 1.5-fold morphological stage. Other landmarks, such as bean or 2-fold, have also been used, but we found the 1.5-fold stage to be the easiest to time unambiguously. The ratio *k* between these two durations (4E/8E and 1.5-fold) was then calculated.

**Figure 2.**
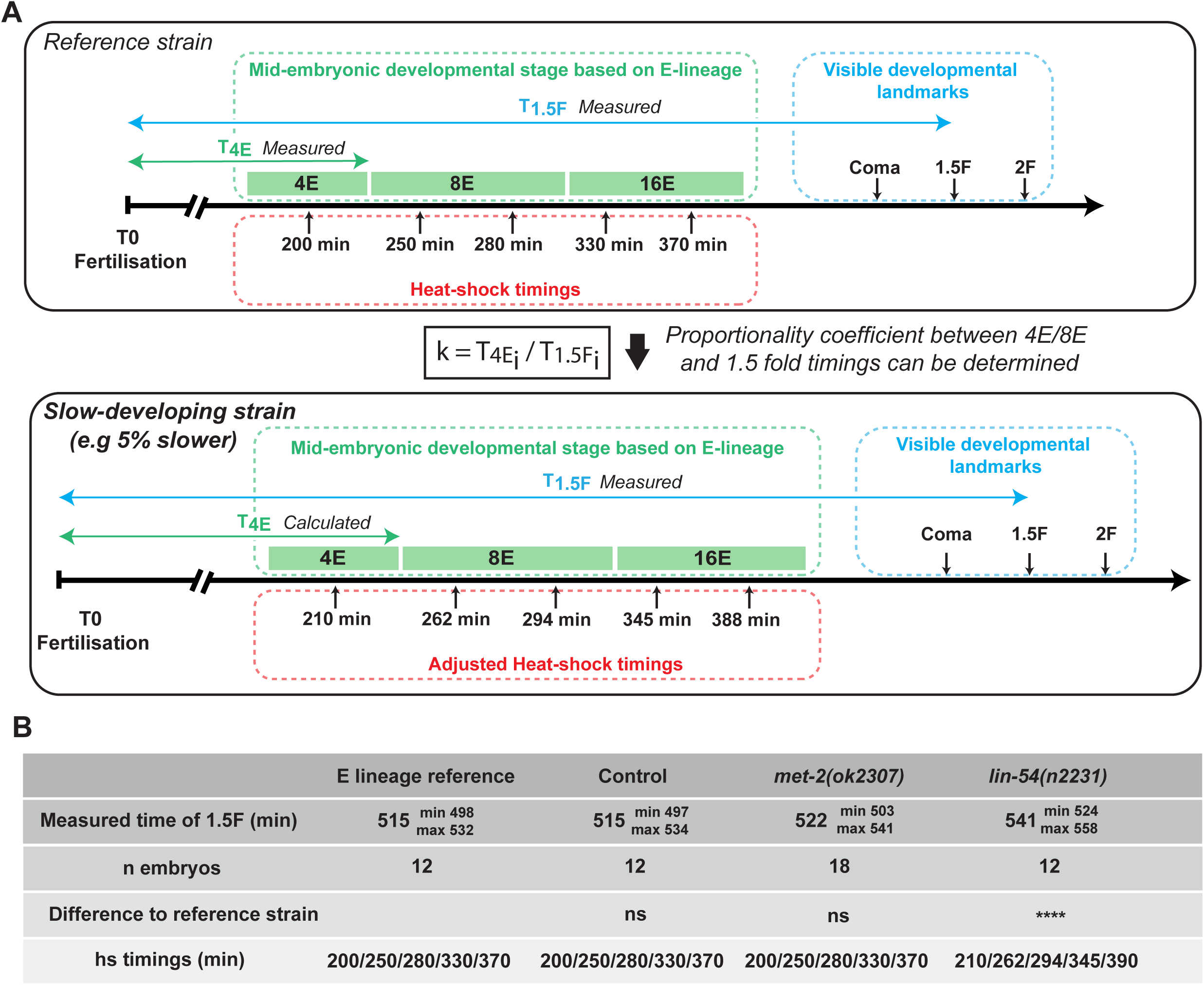
Timing corrections applied to strains with different embryonic developmental speeds. **A.** Five reference cell-fate challenge timings (200 / 250 / 280 / 330 / 370min) were chosen to occur during the 4E 8E and 16E embryonic stages. Validation of these timings compared to embryonic stages were performed in a reference strain endogenously expressing a nuclear GFP under the control of *end-1p* promoter, in the E-lineage (*teIs46* integrated array). The time elapsed to reach three visible, late developmental stages, coma, 1.5-fold and 2-fold, were equally determined in this reference strain. *k* is calculated as the ratio between 4E/8E transition and 1.5-fold timings. The time necessary to reach the 1.5-fold stage in the other genotypes used is also determined and k is used to estimate the timing of the 4E/8E transition in a slower (or quicker) genotype, allowing to apply time-correction to the cell-fate challenge timings. **B.** Measured times to reach the 1.5-fold stage for the reference and the experimental strains used, and adjusted heat shock timings. Mann-Whitney tests with Bonferroni corrections for developmental speed comparisons.

For each tested strain, we measured the time from fertilisation to the 1.5-fold stage. If this duration differed from that of the reference strain, ratio *k* and the measured 1.5-fold timing were used to calculate corrected estimates for the 4E/8E transition and the corresponding heat-shock timings (Fig. 2).

Using this approach, we found that our control strain (carrying only the heat-shock–responsive and body-wall muscle reporter transgenes) and *met-2(ok2307)* mutants developed at the same speed as the reference strain. In contrast, and consistent with previous reports (Costello and Petrella 2019) *lin-54(n2231)* mutants developed 5% more slowly. Accordingly, we adjusted the hs timings for this strain from [200 / 250 / 280 / 330 / 370] to [210 / 262 / 294 / 345 / 388] min post-fertilisation (Fig. 2).

### The timing of the heat-shocks does not impact the final number of nuclei in the embryos

To ensure that differences in the number of *myo-3p::GFP*–positive cells across hs timings did not arise from differences in blastomere proliferation after lineage conversion (for example, a blastomere converted at 200 min could give rise to more *myo-3p::GFP*–positive cells than one converted at 370 min simply because it had more time to divide), we quantified the total number of nuclei in embryos heat-shocked at each timepoint. Specifically, we counted total nuclei using the ubiquitous nuclear reporter *pie-1p::mCherry::his-58* (Fig. 3A) in embryos carrying both the *hs::hlh-1* transgene and the body-wall muscle reporter.

**Figure 3.**
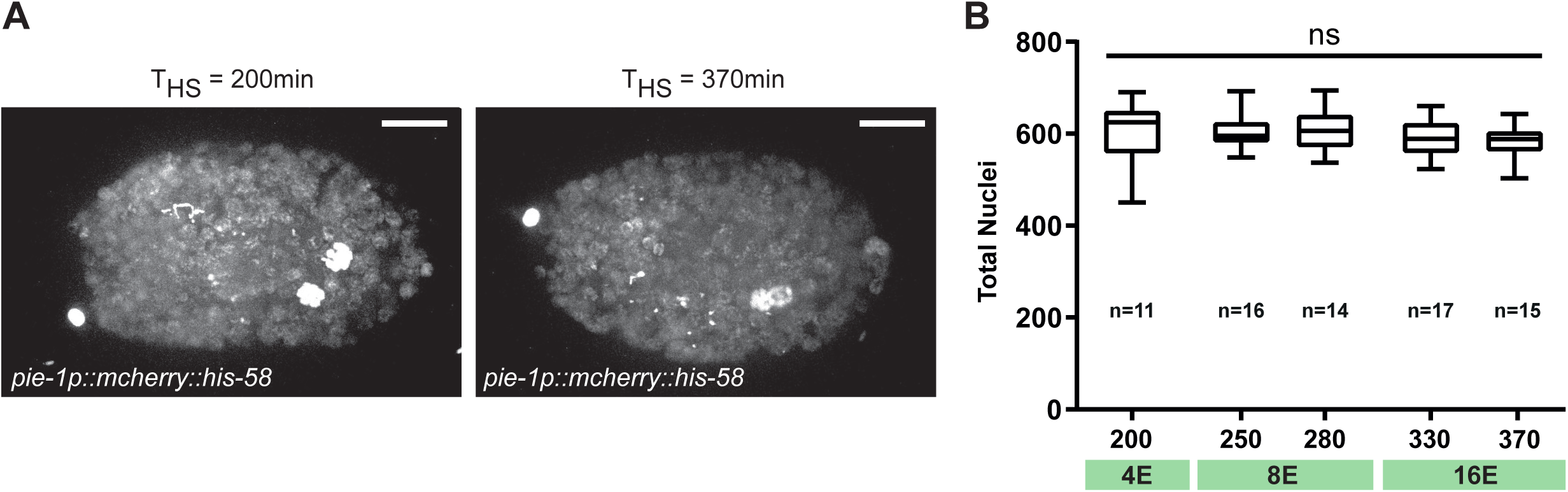
The total number of nuclei after the O/N incubation period is not dependent on the heat-shock timing. **A.** Representative images used for the total nuclei scorings of embryos >12h hours after heat-shock at 200min (4E stage) and 370min (late 16E stage). These embryos carried *hsp-16.41p::hlh-1 cDNA*, *myo-3p::GFP::LacZ::NLS* and *pie-1p::mcherry::his-58* transgenes. Scalebars = 10 µm. **B.** The final total number of nuclei (y axis) is not affected by the timing of the heat-shocks (x axis) in embryos carrying *hsp-16.41p::hlh-1 cDNA* and *myo-3p::GFP::LacZ::NLS* transgenes. Total nuclei were counted >12h after hs based on the expression of the embryonic ubiquitous transgene *pie-1p::mcherry::his-58*. n, number of embryos scored for each condition. Kruskal-Wallis test for rank comparisons.

In these embryos, blastomeres are challenged and converted depending on the hs timing. Therefore, if the timing of the challenge allowed differential proliferation among converted cells, we would expect the total number of nuclei to vary accordingly. However, we found that the total number of nuclei remained stable, at around 600 across all hs timings (Fig. 3B). This value is consistent with the total number of nuclei in *C. elegans* embryos before hatching (558) and with the total number of cells generated during embryogenesis (671) (Sulston et al. 1983). Importantly, these results indicate that, irrespective of when they were performed, the heat-shocks do not influence the proliferation of blastomeres after committing to the BWM fate. Thus, any differences in the number of *myo-3p::GFP*–positive nuclei observed when heat-shocking at different timepoints reflect differences in blastomere plasticity, rather than differences in proliferation after conversion.

### The MCT is a gradual process spanning until the late 16E stage

Analysis of the control *hs::hlh-1* strain showed that the number of converted nuclei consistently and progressively decreases starting from hs at 250 min post-fertilisation (early 8E stage), when almost all nuclei can still be converted, and continues to decline until the 16E stage (hs at 370 min post-fertilisation; Fig. 4A). When heat-shocked at 370 min (late 16E stage), embryos displayed an average of 110 GFP-positive nuclei. Given that *C. elegans* has 95 body-wall muscle cells (Sulston and Horvitz 1977; Sulston et al. 1983), this result indicates that some blastomeres can still be transdetermined into BWM at least until 370 min post-fertilisation.

**Figure 4.**
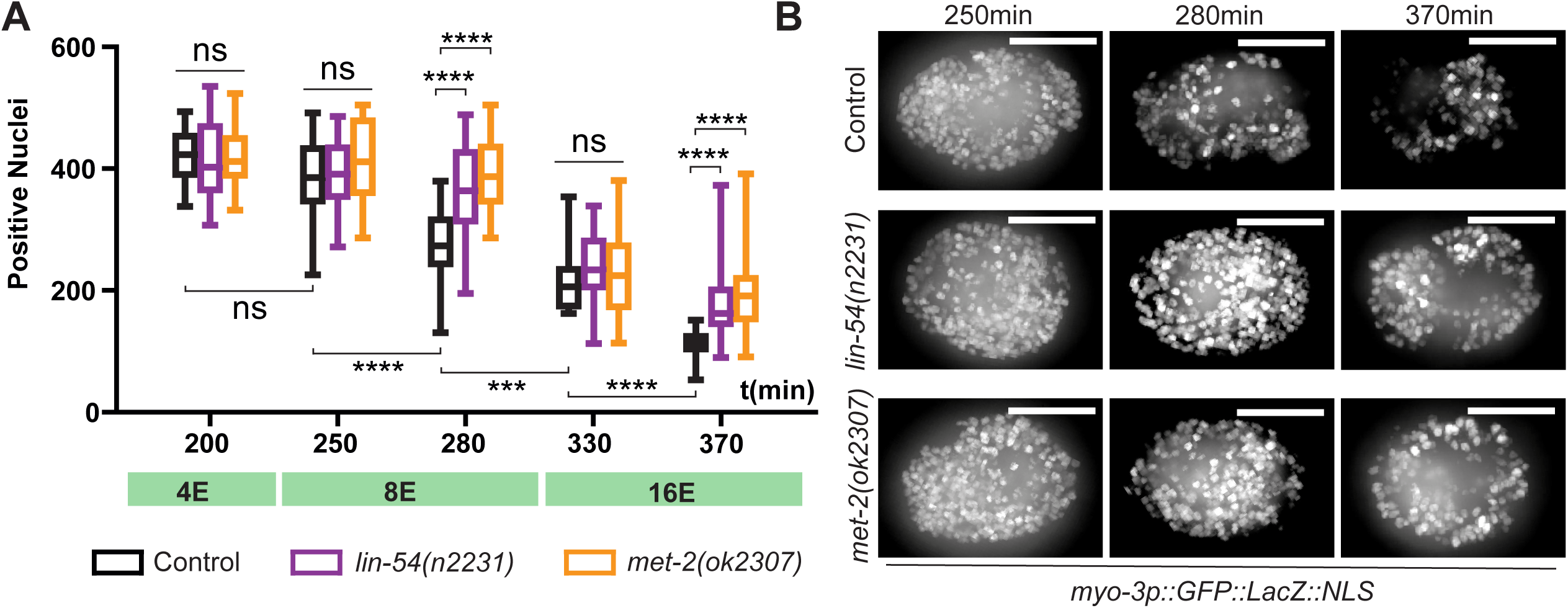
The MCT is a gradual process influenced by the SynMuvBs MET-2 and LIN-54. **A.** Number of *myo-3p::GFP* positive nuclei in control, *lin-54(n2231)* and *met-2(ok2307)* embryos >12h after heatshock. The use of additional precise hs time points shows that the MCT occurs gradually from the early 8E stage rather than sharply between the 8E and 16E time points. More blastomeres can be converted in the SynMuvB mutants *lin-54(n2231)* and *met-2(ok2307)* when heat-shocked at 280min (late 8E stage) or 370min (late 16E) stage highlighting a role for these two factors in promoting commitment. Number of embryos scored at the respective (200/250/280/330/370) timings for each condition: Control n=19/21/25/20/25 – *lin-54(n2231)* n=15/28/27/24/25 – *met-2(ok2307)* n=15/16/24/20/21. Kruskal–Wallis tests and Dunn’s post-hoc tests were used for both gradual MCT in control embryos analysis, and genotype comparison at each timepoint. **B.** Representative images of control (top), *lin-54(n2231)* (middle) and *met-2(ok2307)* (bottom panel) embryos that were heat shocked at 250min, 280min and 370min. Scalebars = 10 µm.

Previous studies documented a sharp and global decline in developmental plasticity (250-280min post-fertilisation). Altogether, our findings refine these timings and demonstrate a progressive reduction at least until the late 16E stage (370 min post-fertilisation).

These observations were made possible by quantifying individual nuclei in challenged embryos, rather than relying on an overall on/off response, and by performing precise and additional later hs timings (200 / 250 / 280 / 330 / 370 min post-fertilisation).

### The DREAM/MuvB complex restricts blastomere multipotency

We recently found that the entire SynMuvB class of genes acted as a brake for Y-to-PDA transdifferentiation (Becker et al. 2026). This class is composed of various chromatin remodelling complexes and heterochromatin-associated proteins (reviewed in (Fay and Yochem 2007)) and was notably identified through its role in repressing *lin-3* during Vulval Precursor Cells specification. Among the SynMuvBs, MET-2 was previously reported to restrict blastomere plasticity at 8E stage, as twice more nuclei still respond to the heat shock challenge in a null mutant (Mutlu et al. 2019). Thus, we wondered whether other members of the SynMuvB class, in particular the DREAM complex, would equally be involved. Specifically, we tested if the MCT would be delayed in DREAM/MuvB core component mutant *lin-54(n2231)*, using *met-2(ok2307)* mutants as positive control.

In line with our hypothesis, we found that blastomeres could still be converted in later *lin-54(n2231)* embryos than in WT, demonstrating extended plasticity. We also found that his was true for *met-2* mutant until the 16E stage (Fig. 4A). Specifically, the number of converted nuclei in these mutants was comparable to the control strain at the earliest hs timings (200 and 250 min post-fertilisation) as well as at 330 min. However, while the number of converted nuclei gradually declines from 250 to 370 min in control embryos, at 280 and 370 min (late 8E and late 16E, respectively), significantly more blastomeres could be converted in *met-2(ok2307)* and *lin-54(n2231)* mutants (Fig. 4A,B).

Together, these findings suggest that the DREAM/MuvB complex is required for blastomere commitment and extended previous reports on MET-2 timely requirement during the MCT (Mutlu et al. 2019). Furthermore, the similar timing and magnitude of *met-2(ok2307*) and *lin-54(n2231)* phenotypes suggest that the MCT is another context where MET-2 and the DREAM/MuvB cooperate (Becker et al. 2026).

## Discussion

### A high-resolution assay to quantify blastomere plasticity

Our approach introduces a high-sensitivity cell-fate challenge assay that differs in several important ways from previous methods used to probe developmental plasticity in *C. elegans*. Notably, instead of relying on a categorical whole-embryo readout—where embryos are scored as either “positive” or “negative”—we quantify individual nuclei that adopt the challenged fate. This provides a much more granular and quantitative assessment of plasticity across developmental time. Conceptually, this brings our method closer to the strategy first used by Gilleard and McGhee (Gilleard and McGhee 2001) who also relied on strong nuclear signals to score transdetermined cells, but with substantially improved resolution and broader applicability. Intriguingly, we observed that the final number of nuclei made, irrespective of when *hlh-1* was over-expressed, is remarkably stable. It could suggest that the number of divisions is independent of the final fate of the cells or, as several cell divisions are observed in WT embryos between the initial endogenous *hlh-1* burst of expression in BWM precursors and terminal differentiation (Guan et al. 2020), reflect the endogenous muscle differentiation programme pace.

A second key improvement concerns the precision of hs timing. By manually selecting 2-cell embryos and mounting them on glass multiwell slides that can be rapidly transferred between temperature-controlled water baths, we achieve near-instantaneous temperature shifts. This allows us to challenge embryos at tightly defined developmental stages, minimizing the temporal variability intrinsic to plate-based or bulk hs approaches. In addition, we leveraged this improved control of the hs by performing several additional hs time points compared to previous studies, increasing temporal granularity.

We further implemented a developmental speed correction to account for genotypes with slower embryogenesis, such as *lin-54(n2231)* mutants. Although slow-growth mutants do not necessarily exhibit a uniform delay across all stages of embryogenesis (Wong et al. 1995; Vajo et al. 1999; Costello and Petrella 2019), the correction we applied proved sufficient. Indeed, *lin-54(n2231)* have a similar MCT-delaying effect with our corrections than *met-2(ok2307)* mutants (for which we did not detect any slow-growth), previously shown to extend the window of blastomere multipotency to the 8E stage (Mutlu et al. 2019). If required, this correction could be further refined in the future by directly integrating the E-lineage timing reporter *teIs46* into slow-growth backgrounds. Since our method allows for imaging just before hs, this would allow precise measurement of the 4E/8E transition in each genetic context. Nevertheless, the current implementation already provides a robust and sensitive method to detect subtle shifts in commitment timing across genotypes.

### Late persistence of multipotency

A long-standing view in the field is that the multipotency-to-commitment transition (MCT) is essentially complete by the 8E stage (Fukushige and Krause 2005) Early work using heat-shock-driven reprogramming assays or lineage-based observations suggested that blastomeres lose their ability to adopt alternative fates after this period (Sulston et al. 1983; Horner et al. 1998; Gilleard and McGhee 2001; Fukushige and Krause 2005; Yuzyuk et al. 2009; Richard et al. 2011; Djabrayan et al. 2012; Spickard et al. 2018; Coraggio et al. 2019; Rothman and Jarriault 2019). Even in *met-2* mutants in which the window of developmental plasticity is prolonged, (Mutlu et al. 2019) previous works reported that global reprogramming has been reported to be ineffective after the 8E stage.

Our quantitative assay with its temporal resolution refines this view by revealing that the MCT is more gradual than previously appreciated and does not conclude at the 8E/16E boundary. Instead, we find that some blastomeres retain a detectable degree of plasticity until the late 16E stage, around 370 minutes post-fertilisation. At this time point, we still observe a substantial number of nuclei adopting BWM identity, despite being well past the developmental stages previously thought to represent the end of competence. It is possible that variations in MCT timing could be observed due to reprogramming transcription factor intrinsic properties (Gilleard and McGhee 2001) or to purely technical explanations: strong or persistent induction from the HLH-1 hs array may enable reprogramming at stages where other arrays fail (eg, differences in promoter strength, array structure, genomic context, induction kinetics or protein persistence). However, previous studies have compared side-by-side *hlh-1* and *end-1* transdetermining abilities and found no difference (Kiefer et al. 2007; Yuzyuk et al. 2009).

Overall, these findings support a model in which the MCT proceeds progressively rather than as a sharp developmental switch, and they highlight the importance of high-resolution assays for accurately capturing the kinetics of blastomere commitment.

### Chromatin reconfiguration as a central driver of blastomere commitment

Our findings further supports the central role of chromatin regulation in controlling the timing of blastomere commitment. LIN-54 and MET-2 are well-established chromatin regulators with roles in restricting gene expression. LIN-54 is a core component of the DREAM/MuvB complex, a major transcriptional repressor complex that coordinates chromatin state with cell-cycle progression and developmental gene regulation (Hoareau et al. 2024). MET-2 is the principal H3K9 histone methyltransferase in early embryos. These findings are also consistent with our recent observations that the nuclear SynMuvA factor LIN-15A is required for a timely MCT (Becker et al. 2026).

The similar timing and magnitude of the phenotypes that we observe in *lin-54(n2231)*, *met-2(ok2307)* and *lin-15A(n767)* mutants (this study, (Becker et al. 2026))—all exhibiting increased plasticity specifically at late 8E and late 16E stages—suggest that these factors may act cooperatively in this process. This interpretation is consistent with their coordinated activity in multiple developmental contexts as SynMuv factors: they function together in vulval precursor cell (VPC) specification, and are all involved during Y-to-PDA transdifferentiation (Becker et al. 2026). Our observations align with earlier work on MES-2/PRC2, which deposits H3K27me3 and is essential for the timely loss of developmental plasticity (Yuzyuk et al. 2009). Loss of MES-2 delays the MCT (similar to the effects we observe for MET-2, LIN-15A and LIN-54 losses) and has been postulated to preserve a permissive chromatin state. Together with recent findings on the H3.3/H3 histone-variant transition (Gleason et al. 2023), these results reinforce a model where multiple layers of chromatin reconfiguration—including H3K27me3, H3K9 methylation, and histone-variant remodelling—act in concert to progressively restrict multipotency and promote irreversible blastomere commitment.

### Potential lineage-dependent differences in the timing of the MCT

Interestingly, a restricted number of blastomeres can still be transdetermined at late 16E stage, suggesting that loss of plasticity is asynchronous across the embryo. Some cells may be more responsive to a given transdetermination TF, and/or retain plasticity for longer. The timing of commitment could reflect the diversity of fates within each lineage. Lineages that produce BWMs remain multifated well past 370 minutes, also generating coelomocytes or gland cells (Sulston et al. 1983). Cells within these lineages may remain more permissive to BWM fate induction by HLH-1, potentially explaining the persistence of transdetermination at late 16E. Consistent with this model, we found that in embryos heat-shocked at 370 minutes, all HLH-1–converted nuclei were positioned at the outer surface of the embryo (Fig. 1B, 4C), spatially close to the natural BWM-producing lineage.

Interestingly, late induction of CHE-1 can still reprogram specific blastomeres (Patel and Hobert 2017; Gleason et al. 2023). Although these studies did not trace the precise lineage origin of each converted cell, neurons arise from multiple highly diversified lineages during embryogenesis (Sulston et al. 1983), opening the possibility that late plasticity in these assays may reflect the prolonged diversity of the corresponding lineages. Taken together, these observations support a model in which blastomere commitment does not occur synchronously across the embryo, but instead may occur at lineage-specific timing – possibly correlated to each lineage fate diversity. In this view, rather than a sharp global transition completed at the 8E/16E boundary, the MCT would be a gradual, lineage-influenced process that closes at slightly different times depending on each lineage.

## Data availability

All data supporting the findings of this study are available within the article and its associated files. Strains and plasmids are available upon request.

## Acknowledgments

Some strains were provided by the CGC, which is funded by NIH Office of Research Infrastructure Programs (P40 OD010440. We thank members of the Jarriault team for critical discussions and constant support.

## Methods

### *C. elegans* culture and general experimental procedures

*Caenorhabditis elegans* strains (see the Reagents Table for a list) were maintained as previously described (Brenner 1974; Daniele et al. 2025) at 20°C unless otherwise specified using OP50 on Nematode Growth Media (NGM) agar plates. Strains were genotyped according to (Morin et al. 2020).

### Statistical analyses

Statistical analyses were performed with PRISM GraphPad 10. For all experiments, the statistical tests used are indicated in the corresponding figure legend. ns: p*>*.0.05, *: p*<*0.05, **: p*<*0.01, ***: p*<*0.001, ****: p<0.0001. See SI Table 1 for all raw numbers and scorings presented in this study.

### Cell fate challenge assays

The detailed protocol can be found on io.protocols (https://www.protocols.io/private/FF795725CC7011F19CC60A58A9FEAC02). Precise synchronisation of the embryos was performed by dissecting gravid mother in M9 with sterile needles and selecting of 2-cell stages embryos (40-75 min post-fertilisation) using a mouth pipette. Selected embryos were placed in individual wells of multiwalled microscopy slides (Paul Marienfeld GmbH & Co. KG 10-well slides Cat. 1216521) with 8 µL of M9 and covered with appropriate coverslips. Slides were sealed using melted Valap (a mixture of 1/3 Vaseline, 1/3 paraffin, 1/3 lanolin). Valap cools down and solidifies almost instantly. Slides were placed in a 20°C water bath until the animals reach the desired timing. At the desired time point, slides were transferred to a 34°C water bath for 30 minutes, and then back in the 20°C water bath overnight for the embryos to develop and lineages to be converted. The hs procedure did not induce any lethality and embryos continued to develop. However, they stopped to develop before morphogenesis due to the lineage conversions and never elongated. Embryos were imaged on a Yokogawa CSU-W1 Leica DMi8 spinning disk microscope using a 0.3 µm z-step. 3D reconstruction from the acquired Z-stack were performed using Imaris 9.8. Nuclei scorings were performed after segmenting the 3D stacks with the surface contour tool. Mitochondrial GFP signal was much weaker and did not interfere with nuclei segmentation. Control scorings of total nuclei were performed following the same protocol, by segmenting nuclei using a *pie-1p::mcherry::his-58* integrated array, driving the expression of the mCherry in all embryonic nuclei.

### Materials

see Reagents Table

